# C1QA is an invariant biomarker for tissue macrophages

**DOI:** 10.1101/2024.01.26.577475

**Authors:** Amir Horowitz, Haocheng Yu, Sonalisa Pandey, Bud Mishra, Debashis Sahoo

**Affiliations:** Marc and Jennifer Lipschultz Precision Immunology Institute, Icahn School of Medicine at Mount Sinai, New York, NY, USA; The Tisch Cancer Institute, Icahn School of Medicine at Mount Sinai, New York, NY, USA; Department of Oncological Sciences, Icahn School of Medicine at Mount Sinai, New York, NY, USA; Department of Pediatrics, University of California San Diego; Departments of Computer Science, Mathematics and Cell Biology, Courant Institute and NYU School of Medicine, New York University, New York City, NY, United States; Department of Computer Science and Engineering, Jacob’s School of Engineering, University of California San Diego

## Abstract

Macrophages play a pivotal role in immune responses, particularly in the context of combating microbial threats within tissues. The identification of reliable biomarkers associated with macrophage function is essential for understanding their diverse roles in host defense. This study investigates the potential of C1QA as an invariant biomarker for tissue macrophages, focusing on its correlation with the anti-microbial pathway. C1QA, a component of the complement system, has been previously implicated in various immune functions. Our research delves into the specific association of C1QA with tissue-resident macrophages and its implications in the context of anti-microbial responses. Through comprehensive systems biology and Boolean analysis of gene expression, we aim to establish C1QA as a consistent and reliable marker for identifying tissue macrophages. Furthermore, we explore the functional significance of C1QA in the anti-microbial pathway. This research seeks to provide valuable insights into the molecular mechanisms underlying the anti-microbial functions of tissue macrophages, with C1QA emerging as a potential key player in this intricate regulatory network. Understanding the relationship between C1QA, tissue macrophages, and the anti-microbial pathway could pave the way for the development of targeted therapeutic strategies aimed at enhancing the host’s ability to combat infections. Ultimately, our findings contribute to the expanding knowledge of macrophage biology and may have implications for the diagnosis and treatment of infectious diseases.

**One Sentence Summary:** C1QA is a fundamental biomarker of tissue macrophages

## Introduction

Macrophages are integral components of the immune system, orchestrating a diverse array of functions crucial for host defense against microbial threats. The identification and characterization of reliable biomarkers associated with macrophage activation and function are essential for a comprehensive understanding of their roles in maintaining tissue homeostasis and combating infections. Among the numerous molecules involved in immune responses, C1QA, a component of the complement system, has garnered attention for its potential as an invariant biomarker specifically linked to tissue-resident macrophages.

The complement system, a fundamental part of the innate immune system, plays a pivotal role in recognizing and eliminating pathogens. C1QA, as part of the C1 complex, has been implicated in various immune processes, yet its specific association with tissue macrophages remains a subject of investigation. This study aims to unravel the potential of C1QA as a consistent and reliable biomarker for identifying and characterizing tissue macrophages, with a particular focus on its relevance in the context of the anti-microbial pathway.

As macrophages exhibit remarkable phenotypic and functional heterogeneity depending on their tissue microenvironment, establishing a robust and invariant biomarker is critical for accurately identifying and studying these immune cells across diverse physiological and pathological conditions. The present research seeks to fill this gap by examining the expression patterns of C1QA in various tissues and conditions, aiming to validate its candidacy as a universal marker for tissue-resident macrophages.

Moreover, beyond its use as a biomarker, we explore the functional significance of C1QA in the anti-microbial responses of tissue macrophages. Through systems biology, Boolean analysis of gene expression data, we aim to elucidate the role of C1QA in modulating macrophage activities against microbial challenges. Unraveling the molecular mechanisms underlying the association between C1QA, tissue macrophages, and the anti-microbial pathway holds promise for advancing our understanding of macrophage biology and may have therapeutic implications for infectious diseases. This investigation thus contributes to the broader landscape of immunological research, offering new insights into the intricate interplay between macrophages and the complement system.

## Materials and Methods

### Data collection and annotation

Publicly available microarray and RNASeq databases were downloaded from the National Center for Biotechnology Information (NCBI) Gene Expression Omnibus (GEO) website. Gene expression summarization was performed by normalizing Affymetrix platforms by RMA (Robust Multichip Average) and RNASeq platforms by normalized counts. We used log2(normalized counts + 1) as the final gene expression value for analyses.

### Computational Approaches

#### StepMiner Analysis

StepMiner is a computational tool that identifies step-wise transitions in a time-series data ^1^. StepMiner performs an adaptive regression scheme to identify the best possible step up or down based on sum-of-square errors. The steps are placed between time points at the sharpest change between low expression and high expression levels, which gives insight into the timing of the gene expression-switching event. To fit a step function, the algorithm evaluates all possible step positions, and for each position, it computes the average of the values on both sides of the step for the constant segments. An adaptive regression scheme is used that chooses the step positions that minimize the square error with the fitted data. Finally, a regression test statistic is computed as follows:

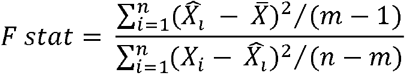

Where *xi* for*i* =1 to *n* are the values, 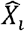 for *i*= 1 to *n* are fitted values. m is the degrees of freedom used for the adaptive regression analysis. 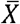 is the average of all the values: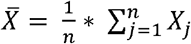For a step position at k,the fitted values 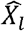 are computed by using 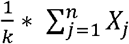 for *i* = 1 to *k* and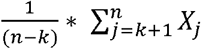 for *i*=*k+*1to *n*

### Boolean Analysis

**Boolean logic** is a simple mathematic relationship of two values, i.e., high/low, 1/0, or positive/negative. The Boolean analysis of gene expression data requires the conversion of expression levels into two possible values. The ***StepMiner*** algorithm is reused to perform Boolean analysis of gene expression data ^2^. **The Boolean analysis** is a statistical approach which creates binary logical inferences that explain the relationships between phenomena. Boolean analysis is performed to determine the relationship between the expression levels of pairs of genes. The ***StepMiner*** algorithm is applied to gene expression levels to convert them into Boolean values (high and low). In this algorithm, first the expression values are sorted from low to high and a rising step function is fitted to the series to identify the threshold. Middle of the step is used as the StepMiner threshold. This threshold is used to convert gene expression values into Boolean values. A noise margin of 2-fold change is applied around the threshold to determine intermediate values, and these values are ignored during Boolean analysis. In a scatter plot, there are four possible quadrants based on Boolean values: (low, low), (low, high), (high, low), (high, high). A Boolean implication relationship is observed if any one of the four possible quadrants or two diagonally opposite quadrants are sparsely populated. Based on this rule, there are six kinds of Boolean implication relationships. Two of them are symmetric: equivalent (corresponding to the positively correlated genes), opposite (corresponding to the highly negatively correlated genes). Four of the Boolean relationships are asymmetric, and each corresponds to one sparse quadrant: (low => low), (high => low), (low => high), (high => high). BooleanNet statistics is used to assess the sparsity of a quadrant and the significance of the Boolean implication relationships ^2, 3^. Given a pair of genes A and B, four quadrants are identified by using the StepMiner thresholds on A and B by ignoring the Intermediate values defined by the noise margin of 2 fold change (+/-0.5 around StepMiner threshold). Number of samples in each quadrant are defined as a_00_, a_01_, a_10_, and a_11_ which is different from X in the previous equation of F stat. Total number of samples where gene expression values for A and B are low is computed using the following equations.

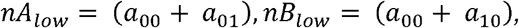

Total number of samples considered is computed using following equation.

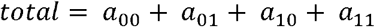

Expected number of samples in each quadrant is computed by assuming independence between A and B. For example, expected number of samples in the bottom left quadrant e_00_ =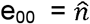 is computed as probability of A low ((a_00_ + a_01_)/total) multiplied by probability of B low ((a_00_+a_10_)/total) multiplied by total number of samples.

Following equation is used to compute the expected number of samples.

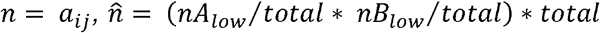

To check whether a quadrant is sparse, a statistical test for (e_00_ > a_00_) or 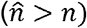is performed by computing S_00_ and p_00_ using following equations. A quadrant is considered sparse if S_00_ is high 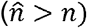 and p_00_ is small.

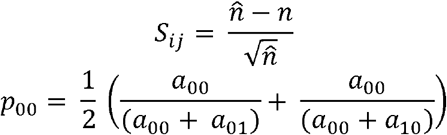

A suitable threshold is chosen for S_00_ > sThr and p_00_ < pThr to check sparse quadrant. A Boolean implication relationship is identified when a sparse quadrant is discovered using following equation.

*Boolean Implication* = (*S*_*ij*_ > sThr, *p*_*ij*_ < pThr)

A relationship is called Boolean equivalent if top-left and bottom-right quadrants are sparse.

*Equivalent* =(*s*_01_ > *sThr, P*01 < *pThr*,s_10_ > *sThr,P*_10_ < *plhr*)

Boolean opposite relationships have sparse top-right (a_11_) and bottom-left (a_00_) quadrants.

*Opposite* = (*s*_00_ > *slhr, P*_00_ < *pThr,s*_11_ > slhr,P_11_ < *pThr*)

Boolean equivalent and opposite are symmetric relationship because the relationship from A to B is same as from B to A. Asymmetric relationship forms when there is only one quadrant sparse (A low => B low: top-left; A low => B high: bottom-left; A high=> B high: bottom-right; A high => B low: top-right). These relationships are asymmetric because the relationship from A to B is different from B to A. For example, A low => B low and B low => A low are two different relationships.

A low => B high is discovered if the bottom-left (a_00_) quadrant is sparse and this relationship satisfies following conditions.

*A low => B high* = (*s*_00_ > *sThr, P*_00_ < *pThr*)

Similarly, A low => B low is identified if the top-left (a_01_) quadrant is sparse.

*A low => B low* = (*s*_01_ > slhr, *P*_01_ < *pThr*)

A high => B high Boolean implication is established if the bottom-right (a_10_) quadrant is sparse as described below.

*A high => B high* = (*s*_10_ > slhr, *P*_10_ < *pThr*)

Boolean implication A high => B low is found if the top-right (a_11_) quadrant is sparse using following equation.

*A high => B low* = (*s*_11_ > *sThr, P*_11_ < *pThr*)

For each quadrant a statistic S_ij_ and an error rate p_ij_ is computed. S_ij_ > sThr and p_ij_ < pThr are the thresholds used on the BooleanNet statistics to identify Boolean implication relationships.

Boolean analyses use a threshold of sThr = 3 and pThr = 0.1. False discovery rate is computed for these thresholds (FDR < 0.000001) by using randomly permuting gene expression data.

### SMaRT analysis

The Signatures of Macrophage Reactivity and Tolerance (SMaRT) offer a comprehensive quantitative and qualitative framework for evaluating macrophage polarization across various tissues and conditions. This study unveils a gene signature remarkably conserved across diverse tissues and conditions, comprising a set of 338 genes derived from a Boolean Implication Network model of macrophages. This model effectively identifies macrophage polarization states at the single-cell level, encompassing a spectrum of physiological, tissue-specific, and disease contexts. Remarkably, this signature demonstrates robust associations with outcomes in several diseases, underscoring its potential as a valuable predictive tool.

The algorithm uses three clusters C#13, C#14, C#3 from the published macrophage network and uses composite scores of C#13, C#14-3, and C#13-14-3 to identify macrophage polarization states (See function getCls13, getCls14a3, getCls13a14a3, and order Data in github codebase BoNE/SMaRT/MacUtils.py and the outputs in BoNE/SMaRT/macrophage.ipynb). To compute the composite score, first the genes present in each cluster were normalized and averaged. Gene expression values were normalized according to a modified Z-score approach centered around StepMiner threshold (formula = (expr – SThr – 0.5)/3∗stddev). Weighted linear combination of the averages from the clusters of a Boolean path was used to create a score for each sample. The weights along the path either monotonically increased or decreased to make the sample order consistent with the logical order based on Boolean Implication relationships. The samples were ordered based on the final weighted (−1 for C#13, 1 for C#14 and 2 for C#3) and linearly combined score. Performance is measured by computing ROC-AUC. Bar plots show the ranking order of different sample types based on the composite scores of C#13, path C#14-3, or C#13-14-3. Violin plots shows the distribution of scores in different groups. P values are computed with Welch’s Two Sample t-test (unpaired, unequal variance (equal_var = False), and unequal sample size) parameters.

### Statistical analyses

Gene signature is used to classify sample categories and the performance of the multi-class classification is measured by ROC-AUC (Receiver Operating Characteristics Area Under The Curve) values. A color-coded bar plot is combined with a density or violin + swarm plot to visualize the gene signature-based classification. All statistical tests were performed using R version 3.2.3 (2015-12-10). Standard t-tests were performed using python scipy.stats.ttest_ind package (version 0.19.0) with Welch’s Two Sample t-test (unpaired, unequal variance (equal_var = False), and unequal sample size) parameters. Multiple hypothesis corrections were performed by adjusting p values with statsmodels.stats.multitest.multipletests (fdr_bh: Benjamini/Hochberg principles). The results were independently validated with R statistical software (R version 3.6.1; 2019-07-05). Pathway analysis of gene lists were carried out via the Reactome database and algorithm.32 Reactome identifies signalling and metabolic molecules and organizes their relations into biological pathways and processes. Kaplan–Meier analysis is performed using lifelines python package version 0.14.6.

## Results

### C1QA is identified as an invariant biomarker of tissue macrophages

Macrophages and the complement system are intricately interconnected components of the immune response and inflammation. It is crucial to comprehend the process by which monocytes in the blood undergo differentiation into macrophages within solid tissues (**Fig 1a**). Utilizing the universally expressed seed gene TYROBP in monocytes and macrophages, we identify candidate genes exclusive to macrophages. This involves identifying genes analogous to TYROBP in solid tissues and ranking them based on correlation (**Fig 1b**). Furthermore, we assess the differential expression between solid tissue and blood to gain insights into this differentiation process (**Fig 1c**). C1Q genes are highly ranked. We discovered a fundamental Boolean implication relationship between TYROBP (universal biomarker of macrophages) and C1QA (part of the complex that binds to antibodies, **Fig 1d**). This relationship was conserved in human, mouse, rat, baboon, monkey, and dog. Datasets were derived from diverse tissue types profiled in human microarray (n = 25,955, GSE119087, Affymetrix U133 Plus 2.0) and mouse microarray (n = 11,758, GSE119085, Affymetrix Mouse 430 2.0), rat microarray (n = 11,599, Pooled GEO, Rat 230 2.0), baboon RNAseq (n = 767, GSE98965), monkey RNASeq (n = 288, GSE219045), and dog RNASeq (n = 300, GSE219045). The tissue types were carefully annotated and categorized into three different states: blood (liquid), solid tissue (colon, lung, brain etc.), and cell line. Detailed analysis in each tissue type revealed that the Boolean implication relationship is tissue specific: C1QA Equivalent TYROBP (solid), C1QA high => TYROBP high (liquid), C1QA low (cell line). Each of these relationships was also conserved across species (**Fig 1e**). Such strong relationships are called *invariant* because the Boolean formula always evaluates to true. The asymmetric invariant Boolean relationships provide a computational platform to define a continuum of biological states. Previously, we used these approaches to characterize B cell differentiation^3, 4^, colon differentiation^5, 6^, inflammatory bowel disease spectrum^7^, macrophage polarization states^8^.

**Figure 1:**
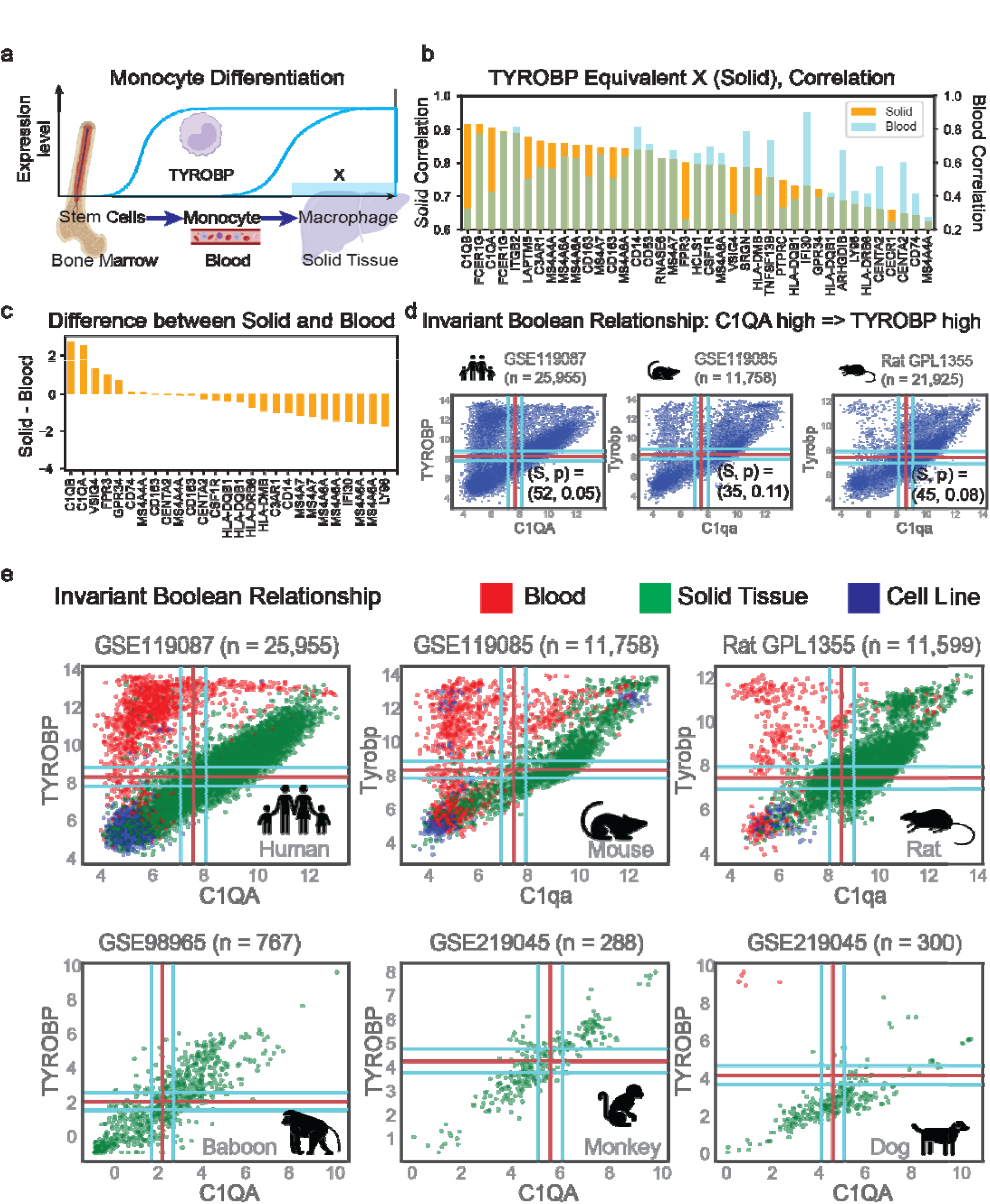
TYROBP and C1QA are associated with tissue specific invariant Boolean Implication relationship. (a) Schematic experimental design to study monocyte to macrophage differentiation. (b) The search for potential candidates meeting the criteria of TYROBP Equivalent X in solid tissue, as annotated in GSE119087, involves ranking them based on their correlation coefficients within solid tissue, concurrently evaluating their correlation coefficients in blood tissue. (c) The gene candidates are ranked based on the difference in mRNA expression between solid and blood tissue, while concurrently satisfying the criteria of TYROBP Equivalent X in solid tissue as annotated in GSE119087. (d) Boolean implication relationship C1QA high => TYROBP high is conserved in human (GSE119087; n = 25,955), mouse (GSE119085; n = 11,758) and rat (Pooled GEO samples from platform GPL1355, n = 21,925) datasets. (e) Scatterplots depicting the relationship between C1QA and TYROBP, with blood samples in red, solid tissue samples in green, and cell-line samples in blue, across various species including human (GSE119087), mouse (GSE119085), rat (pooled GEO samples from platform GPL1355), baboon (GSE98965), monkey (GSE219045), and dog (GSE219045) datasets.

### Boolean analysis reveals a continuum of monocyte to macrophage differentiation

Monocytes, classified as phagocytic cells and integral members of the mononuclear phagocyte system, initially circulate in the bloodstream. Subsequently, they migrate into tissues, where they transform into macrophages. Although there has been extensive research into the differentiation of monocytes into macrophages, identifying the precise intermediate states has proven challenging due to the absence of suitable biomarkers. To address this issue, the MiDReG analysis approach was devised within the context of systems biology. This method seeks to elucidate these intermediate states by capitalizing on the strong symmetric Boolean implication relationship observed between TYROBP and C1QA in solid tissues, as well as the asymmetrical Boolean implication found in blood. These findings have motivated the development of a model for macrophage differentiation centered around the expression of C1QA (**Fig 2a**). The level of C1QA in monocytes is similar to B, T, and NK cells and elevated in macrophages and dendritic cells (GSE46903, n=384, **Fig 2b**). In addition, a sequential increase in C1QA expression is discovered with the three well-defined classes of monocytes derived from the umbilical cord blood (GSE195727, n = 151, 25-41 GA, **Fig 2c**). The differential expression of C1QA is validated in six additional datasets (4 human and 2 mice, **Fig 2d**). Comprehensive analysis of C1QA in mononuclear phagocytes (MNP-VERSE,^9^ GSE178209) demonstrated high expression in diverse solid tissue types (**Fig 2e**). Together, the data suggests that C1QA expression is critical for the differentiation of monocytes into macrophages.

**Figure 2:**
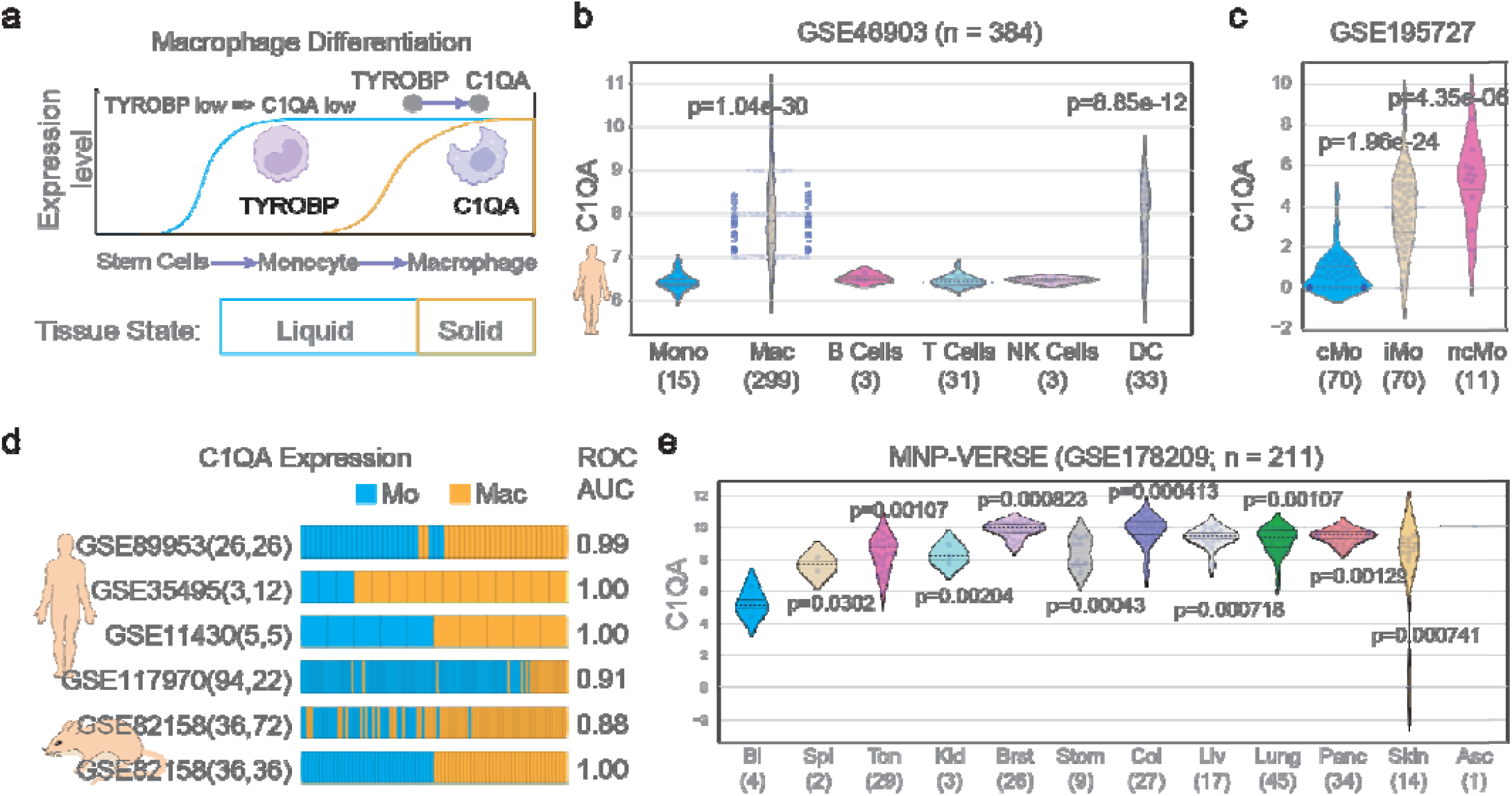
Validation of the Consistent Boolean Model for Monocyte to Macrophage Differentiation. (a) Schematic diagram to study monocyte to macrophage differentiation. (b) C1QA expression pattern in monocyte, macrophages, B cell, T cells, NK cells and DC cells in GSE46903 (n = 384). (c) Expression patterns of C1QA across three different subsets of monocytes in cord blood (GSE195727): classical (n = 70), intermediate (n = 70) and non-classical (n = 11). (d) C1QA expression Is ordered from low to high from left to right highlighting the sample annotation from monocytes and macrophages in four human (GSE89953, GSE35495, GSE11430, GSE117970) and two mouse datasets (GSE82158). (e) C1QA expression patterns in mononuclear phagocytes in diverse tissues (MNP-VERSE, GSE178209, Blood, Spleen, Tonsil, Kidney, Breast, Stomach, Colon, Liver, Lung, Pancreas, Skin, and Ascites).

### Solid tissue CD16 expression is invariantly linked to the negative regulation of classical complement activation and promoting immune tolerance through LAIR1 expression

Classical complement activation is triggered by C1q recognition of IgG, IgM, CRP/Pentraxins. Specific to IgG, human C1q binds to hIgG1 and hIgG3, the same isotypes as human CD16 (*FGCR3A*)^10^. Mouse C1q binds to mIgG2a and mIgG2b, the same as murine CD16 (*FCGR3*). Strikingly, our Boolean network models revealed an invariant dependency of FCGR3A on C1q co-expression (in both mice and humans) (**Fig 3c**), suggesting that C1q is always present to compete with CD16 for IgG in solid tissues. C1q is thought to bind IgG with significantly greater affinity compared to CD16; thus, how would it be possible for Ab-mediated CD16 signaling to ever play out?

**Figure 3:**
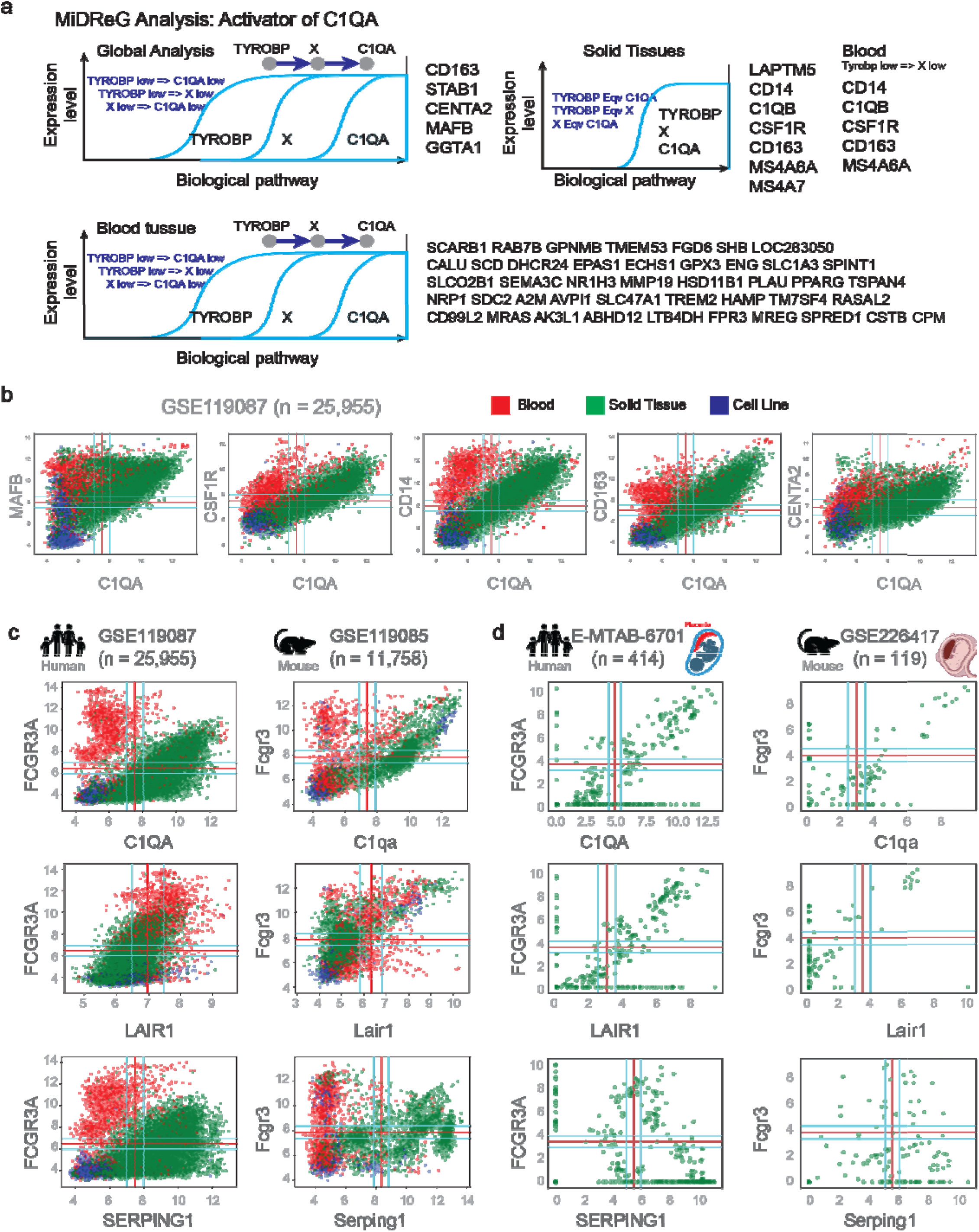
Identification of Activator of C1QA using MiDReG analysis. (a) Study design and identification of activators of C1QA in solid, and blood tissues. (b) Scatterplots of selected genes with C1QA highlighting blood in red, solid in green and cell-line in blue in GSE119087. (c) Scatterplots of genes associated with classical complement activation pathway with CD16 (FCGR3A, Fcgr3) highlighting blood in red, solid in green and cell-line in blue in human (GSE119087) and mouse (GSE119085). (d) Scatterplots of genes associated with classical complement activation pathway with CD16 (FCGR3A, Fcgr3) in human (E-MTAB-6701) and mouse (GSE226417) placenta single cell dataset (pseudo-bulk of different clusters of cells).

The C1-inhibitor (*SERPING1*) competes with the C1r and C1s proteases for binding the C1q molecule, whereby the C1r and C1s proteases are displaced and can no longer cleave complement C2 and C4 proteins to allow formation of C3 convertase^11-13^. As a direct consequence, lack of C3 convertase prevents the formation of C3a, and induction of C3aR-mediated inflammation as well as C3b and induction of CD11b/CD18 (CR3) and CD11c/CD18 (CR4) mediated phagocytosis as well as C5 convertase and formation of the membrane attack complex (MAC) that induces apoptosis. Thus, if *FCGR3A* is expressed in solid tissues (regardless if by NK, myeloid, or granulocyte subsets), C1q may be invariantly inhibited and unable to bind IgG (but also IgM and CRP) and the complement system is shut down. Of great importance, our analyses also revealed that the inhibitory LAIR1 receptor, which recognizes the C1 complex (C1QA/B/C, C1r, C1s)^14^ when bound to antigen (IgG,IgM,CRP), is also invariantly linked to *FCGR3A* expression in solid tissues. To the best of our knowledge, this is the first ever example of a direct inhibitory counterpart to CD16-mediated activation. Stated differently, LAIR1 is inhibitory and can be induced on most immune cells. Thus, if allowed to interact with C1 complex, this would promote inhibition and immune tolerance. Conversely, if CD16 binds IgG instead of C1q, this leads to direct activation, and when expressed by NK cells, CD16 is the only known activating receptor that does not require any co-stimulation.^15^

Additionally, we had access to human and mouse datasets of single-cell RNA sequencing of decidual and placental tissues from early pregnancy where the physical trophoblast barrier separating placenta from decidua is still intact (**Fig 3d**). Data were transformed into a pseudobulk format and we observed near identical relationships between CD16 and C1q, LAIR1 and C1-inhibitor. The data fundamentally indicates that C1q in solid tissues serves a predominantly tolerogenic role, where CD16 regulates and modulates C1q functionality based on CD16 expression abundance affording itself to rapid response against invading pathogens.

### L. monocytogenes infection of human placental Hofbauer cells and E-coli infection of monkey decidual macrophages differently regulates expression of C1q and CD16 but promotes polarization into M1 reactive macrophages

Cumulatively, the data unveil novel insights into the regulatory role of C1q in immune tolerance within solid tissues, and how bloodborne microbial infections influence C1q expression to orchestrate antimicrobial immunity (Fig 4a). If our initial interpretations hold true, the suppression of C1q expression and the complement system in the fetal/placental microenvironment during early pregnancy may confer advantages for fetal development. To test this hypothesis, we examined publicly available microarray and RNA-seq datasets encompassing human peripheral blood from diverse groups, including healthy controls, patients with bacterial infections/sepsis, RSV and influenza patients, and malaria-infected placenta (**Fig 4b-c**). Our findings indicate a general trend of increased C1QA expression and M1 macrophage polarization in response to infections, with sepsis patients and malaria-infected placenta showing significantly elevated C1QA expression—validated across 14 original datasets (**Fig 4d**).

**Figure 4:**
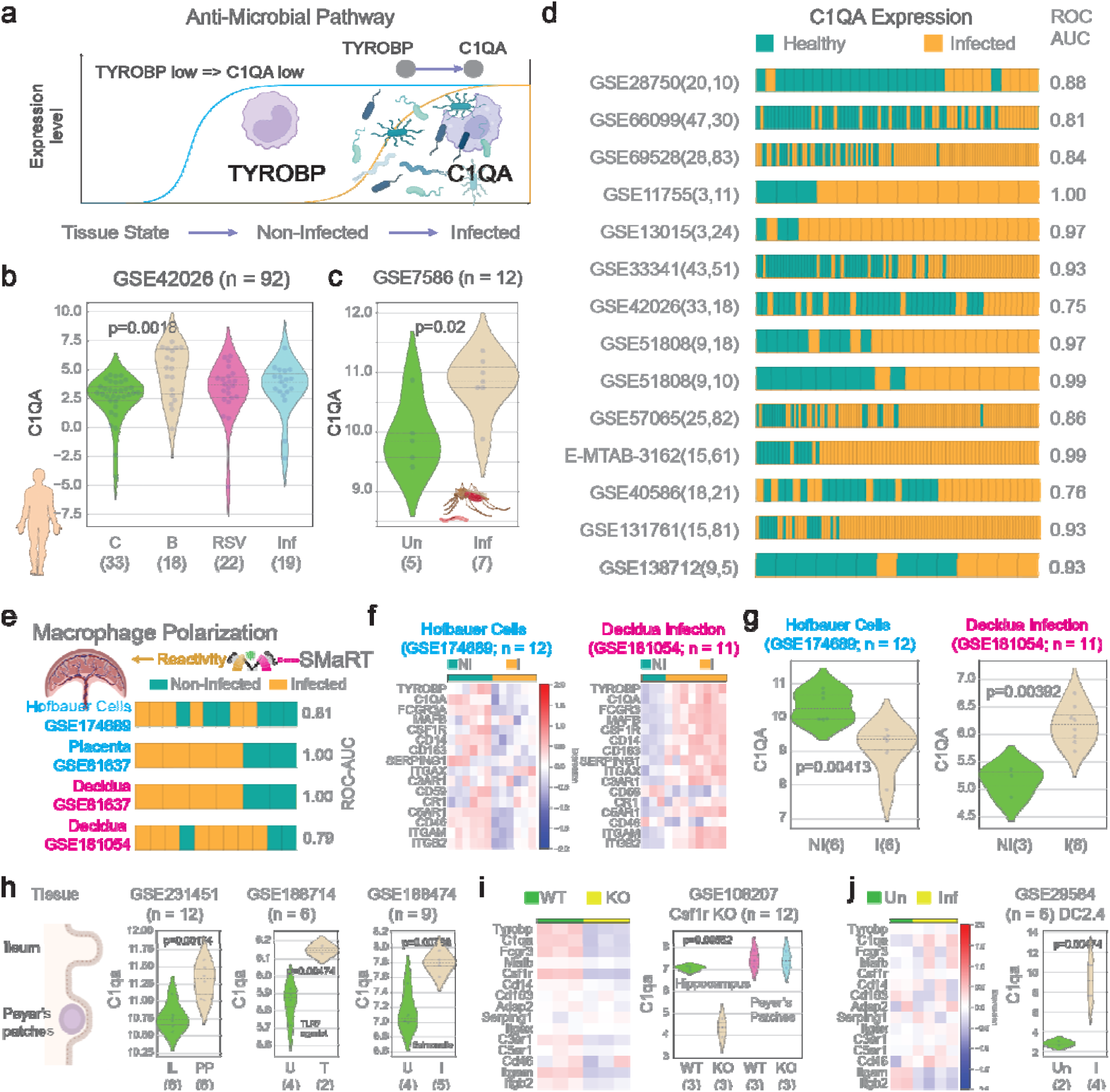
The role of C1QA in anti-microbial pathway. (a) Schematic study of C1QA expression in microbe infection datasets. (b) Violin plots of C1QA expression during bacterial, RSV, and Influenza infection (GSE42026). (c) Violin plots of C1QA expression during placental malaria (GSE7586). (d) Bar plots depicting C1QA expression ordered from low to high, arranged from left to right, with emphasis on sample annotations (Healthy vs Infected). (e) C#13 based SMaRT model of macrophage polarization during infection. (f) Heatmaps of selected genes (C1QA activators, CD16 and the complement pathway) in Hofbauer cells (GSE174689) and Decidua (GSE181054) in non-infected and infected settings. (g) Violin plots of C1QA expression in Hofbauer cells (GSE174689) and Decidua (GSE181054) in non-infected and infected settings. (h) Violin plots of 1QA expression in Peyer’s patches datasets (GSE231451, GSE188714, GSE188474) in uninfected and infected settings. (i) Heatmaps of selected genes (C1QA activators, CD16 and the complement pathway) and Violin plots of C1QA expression in Csf1r knockout hippocampus and Payer’s patches. (j) Heatmaps of selected genes (C1QA activators, CD16 and the complement pathway) and Violin plots of C1QA expression in DC2.4 cell-line (GSE29584, uninfected vs Infected).

RNA sequencing data from publicly available human studies of Listeria infection in cells isolated from healthy pregnant women were used to evaluate how C1q and CD16 are differently modulated if an infection is within the fetal or maternal microenvironment. Hofbauer cells (HBCs) are eosinophilic histiocytes believed to be of macrophage origin, restricted to the placenta, heavily enriched during early pregnancy and most likely involved in the vertical transmission of pathogens from mother to fetus and are highly susceptible to Listeria infection.

Our Boolean network analyses identified five critical genes required for expression of C1q, all of which play important roles in anti-microbial immunity and macrophage polarization (**Fig 4e**). SMaRT predicts pro-inflammatory states in both HBCs and Decidua (**Fig 4e**). Anti-microbial protection from infectious pathogens is critical for successful organogenesis and fetal development. Our data suggests a pro-tolerogenic role for C1q, we hypothesized that C1q expression (and perhaps its driver genes) would be suppressed upon infection.

Indeed, both C1q and CD16 were significantly diminished on infected HBCs along with expression of *CSF1R, MAFB, CD14, and CD163* compared to uninfected controls (**Fig 4f**). Conversely, *E. coli* infection had the opposite effect on monkey decidual macrophages whereby C1q and CD16 are strongly upregulated as well as the driver genes for C1q, suggesting a hierarchy in importance to promote immune tolerance in the decidua, even at the cost of infection, and possibly revealing therapeutic pathways that can be exploited in the clinic (**Fig 4f-g**).

### C1QA is associated with severe sepsis and flu

To check if C1QA expression is relevant for human diseased, we analyzed sepsis (GSE65682, GSE224146) and flu datasets (GSE101702, GSE61821, GSE21802, GSE42641) that are annotated with severity of the disease. C1QA high expression was associated with 28-days mortality in a sepsis study (GSE65682, n = 479, p = 0.032, **Fig 5a**). To study the organism-wide response to sepsis, we analyzed C1QA gene expression across tissues in a model leading to sepsis using cecal ligation and puncture (CLP).^16^ We examined C1QA gene expression changes in 13 tissues including bone marrow, brain, colon, heart, inguinal lymph nodes (iLNs), kidney, liver, lung, peripheral blood mononuclear cells (PBMCs), skin, small intestine, spleen and thymus covering early and late effects—and from untreated control mice (**Fig 5b**). As shown in the figure, C1QA expression consistently upregulated in moderate cases of sepsis from Day 0.5 to Day 1. Whereas in severe cases, the difference is observed only in spleen, thymus and lung samples (**Fig 5b**). Examination of peripheral blood mononuclear cell (PBMC) datasets from individuals with severe influenza revealed an elevated expression of C1QA in severe cases (GSE101702, GSE61821, GSE21802, **Fig 5c-e**). The analysis of C1QA expression encompasses five distinct cell types—Alveolar Macrophages, Lymphocytes, Monocytes, Neutrophils, and epithelial cells—in mice subjected to sublethal and lethal doses of H1N1 (GSE42641, **Fig 5f**). Notably, differential expression of C1QA is observed exclusively within Monocyte populations (**Fig 5f**).

**Figure 5:**
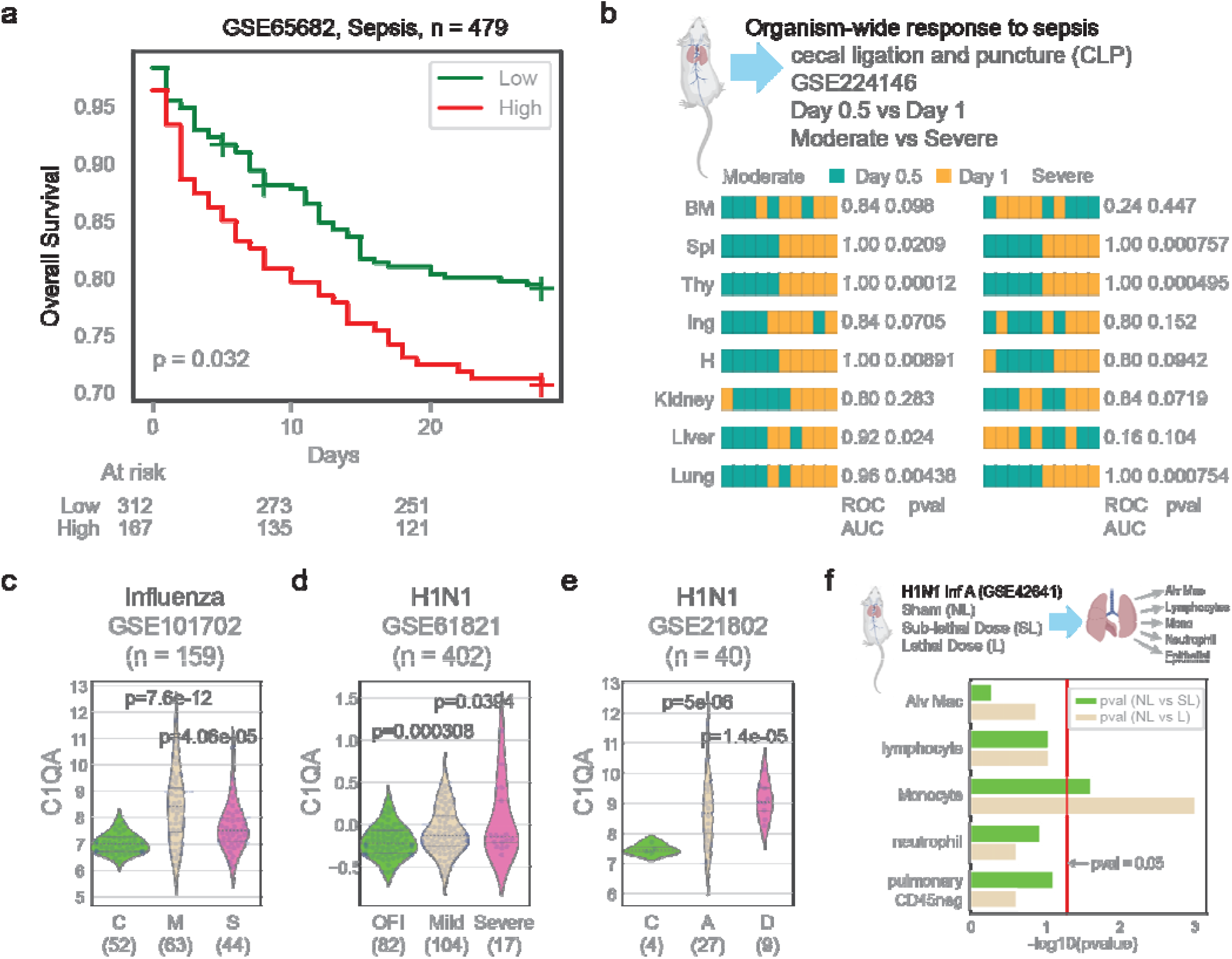
C1QA expression is associated with severe sepsis and flu. (a) C1QA high expression predicts 28-days mortality in sepsis patients (GSE65682). (b) C1QA high expression predicts moderate and severe sepsis in different mouse tissues (GSE224146, Day 0.5 vs Day 1). (c) Violin plots of C1QA expression in healthy (C, n=52), moderate (M, n=63) and severe (S, n=44) influenza (GSE101702). (d) Violin plots of C1QA expression in other febrile illness (OFI, n=82), mild (n=104) and severe (n=17) H1N1 infection (GSE61821). (e) Violin plots of C1QA expression in healthy controls (C, n=4), alive (A, n=27) and dead (D, n=9) H1N1 infected patients (GSE21802). (f) Comparison of C1QA expression in different cell types (Alveolar Macrophages,Lymphocytes, Monocytes, Neutrophils, and Epithelial cells) from mice infected with sub lethal dose and letha of H1N1 infection. Red line refers to the pvalue threshold of 0.05.

## Conclusion

In conclusion, this study sheds light on the potential of C1QA as an invariant biomarker for tissue macrophages and its association with the anti-microbial pathway. Through comprehensive analyses of C1QA expression patterns in various tissues and under different conditions, we have provided evidence supporting its candidacy as a reliable marker for identifying and characterizing tissue-resident macrophages. The establishment of such a marker is crucial for advancing our understanding of macrophage biology, allowing for more accurate and consistent identification across diverse physiological contexts.

Furthermore, our investigation into the functional significance of C1QA in the anti-microbial pathway has uncovered intriguing insights into the role of this complement component in modulating macrophage responses to microbial challenges. The intricate interplay between C1QA and tissue macrophages suggests a potential regulatory mechanism that warrants further exploration. Understanding these molecular mechanisms not only enhances our comprehension of macrophage biology but also opens avenues for developing targeted therapeutic strategies aimed at bolstering the host’s immune defense against infections.

As the field of immunology continues to evolve, the identification of reliable biomarkers and the elucidation of their functional relevance become paramount for advancing diagnostic and therapeutic approaches. C1QA, with its dual role as a potential biomarker and participant in the anti-microbial response, emerges as a key player in this intricate network. The findings presented in this study contribute valuable knowledge to the ongoing discourse on macrophage biology and complement-mediated immunity.

In summary, the exploration of C1QA as a biomarker for tissue macrophages and its involvement in the anti-microbial pathway underscores its significance in the broader landscape of immune responses. Further research and translational efforts in this direction may lead to innovative strategies for diagnosing and treating infectious diseases, ultimately benefiting human health.

## Contributors

Conceptualization: D.S, A.H

Methodology: D.S, A.H, H.Y, B.M

Investigation: D.S, A.H, H.Y, B.M

Visualization: D.S, A.H., H.Y, S.P

Funding acquisition: D.S, A.H, B.M

Project administration: D.S, A.H

Supervision: D.S, A.H

Writing – original draft: D.S, A.H

Writing – review & editing: D.S, A.H, H.Y, S.P, B.M

D.S, and A.H have accessed and verified the underlying data. All authors read and approved the final version of the manuscript.

### Data Sharing

All data are available in the main text or the supplementary materials. The codes are available in https://github.com/sahoo00/BoNE.

## Declaration of interests

SP and DS are co-founders of the company Shanvi. SP is the President of Shanvi. All other authors have no competing interests. The authors declare that they have no financial conflict of interests for this study.

## Acknowledgements

This work was supported by the National Institutes for Health (NIH) grant R01-AI155696 (to DS). Other sources of support include: R01-GM138385 (to DS), and UG3TR003355 (to DS). Images in the figures were created with BioRender.com.

## Notes

### Competing Interest Statement

The authors have declared no competing interest.

